# A conserved Arf-GEF modulates axonal integrity through RAB-35 by altering neuron-epidermal attachment

**DOI:** 10.1101/2025.06.03.657768

**Authors:** Igor Bonacossa-Pereira, Sean Coakley, Massimo A. Hilliard

## Abstract

Neurites of sensory neurons densely innervate the skin and are embedded within it. These delicate structures are exposed to acute and chronic mechanical strain and yet their integrity is maintained throughout life. Although evidence suggests a neuroprotective role for the skin, the molecular pathways involved are still poorly understood. In *C. elegans*, the cytoskeletal molecule UNC-70/β-spectrin functions in synergy with the small GTPase RAB-35 within the skin to stabilize neuron-epidermal attachment structures against mechanical strain and prevent movement-induced damage to mechanosensitive axons. However, the full suite of molecules regulating these specialized attachments remains elusive. Here, through an unbiased forward genetic screen we have identified a guanine nucleotide exchange factor (GEF) previously associated with the endocytic-recycling machinery, AGEF-1a, that impacts axonal maintenance. We show that AGEF-1a functions selectively within the skin to regulate the integrity of the embedded axons. Mechanistically, we reveal that this effect is achieved through an interaction between AGEF-1a and epidermal RAB-35 to facilitate its activation, which in turn modulates the neuron-epidermal attachments. Finally, we demonstrate that the function of this GEF is highly conserved, with the expression of its human ortholog, BIG2 capable of replacing AGEF-1a. Together, we reveal the specific molecular machinery responsible for fine-tuning neuron-epidermal attachments and maintaining axonal integrity during life.

## Introduction

Neurons extend disproportionally long axons that develop specialized attachments with surrounding cells. This is especially evident in sensory axons innervating the epidermis. Neuron-epidermal attachments induce the embedment of sensory axons within the skin, tightly adhering and mechanically coupling these tissues ^1–5^. Due to their slender architecture and anatomical localization, these axons are vulnerable to movement-induced mechanical strain ^1,5–10^.

Spectrins provide mechanical resistance and maintain membrane shape against strain in all cells ^11^. Axons are enriched in β-spectrin ^12–14^, which contributes to axonal elasticity and force resistance ^1,6,10^. We have recently demonstrated, using the nematode *C. elegans* as a model system, that axonal β-spectrin is not sufficient to protect sensory neurons from damage, and instead functions non-cell-autonomously within the epidermis to maintain the integrity of the posterior lateral mechanosensory neuron (PLM) ^5,10^. During development, these neurons attach and embed within the epidermis via specialized neuron-epidermal attachments ^1,15^. These attachments must be mechanically resilient to prevent damage to the PLM axon upon body movement. We have shown that the mechanical resilience of the neuron-epidermal attachments is mediated by β-spectrin and a small GTPase RAB-35, which function together in the epidermis to stabilize these attachments against movement and maintain axonal integrity ^5^.

Small GTPases are molecular switches that cycle between active and inactive states to regulate various cellular pathways, including cytoskeleton remodeling and membrane recycling ^16^. Active GTPases bind GTP, and require GTPase activating proteins (GAPs) to accelerate GTP hydrolysis and cause their subsequent inactivation ^16^. Conversely, inactive GTPases bind GDP, and require guanine nucleotide exchange factors (GEFs) to catalyze the release of GDP and binding of a new GTP molecule to become active ^16^. Chronic activation of the epidermal small GTPase RAB-35, induced either by the loss of the GAP TBC-10 or by a mutation that renders RAB-35 constitutively active, causes catastrophic PLM axon breaks when the epidermal spectrin network is disrupted ^5^. This supports a model where regulation of RAB-35 activity is required, redundantly with epidermal β-spectrin, to maintain axonal integrity. While two RAB-35-specfic GEFs, RME-4 and FCNL-1, have been identified to activate RAB-35 in the context of endocytosis and cell corpse clearance ^17,18^, in the context of axonal maintenance RME-4 is only partially involved and FCNL-1 has no effect ^5^. This suggests the existence of unknown RAB-35 activators that function to regulate axonal maintenance.

To identify novel RAB-35 activators modulating axonal integrity, we performed an unbiased genetic screen for suppressors of axonal damage in sensitized animals where the PLM axon spontaneously breaks. From this screen, we identified AGEF-1a, an Arf GEF previously associated with endocytic-recycling and cell-attachment polarity pathways ^19–22^. We demonstrate that this molecule is evolutionarily conserved, with its human ortholog BIG2 capable of replacing the nematode gene function. Next, we reveal that AGEF-1a functions within the skin, in a non-cell-autonomous fashion, to modulate the integrity of the embedded axon. Finally, we show that AGEF-1a interacts with, and facilitates, RAB-35 activation *in vivo* to impact axonal integrity by altering the neuron-epidermal attachments.

## Results

### AGEF-1a/BIG2 regulates axonal integrity non-cell-autonomously in the epidermis

Small GTPase activation by GEFs is a highly contextual and redundant mechanism, thus predicting the GEFs relevant for a given functional outcome is challenging ^16^. To identify novel GEFs relevant to axonal integrity, we conducted an unbiased forward genetic screen in a sensitized genetic background in which sensory axons spontaneously break due to mechanical strain. Specifically, we used animals expressing a dominant negative *unc-70/β-spectrin* cDNA selectively within the epidermis (*vdSi2 = SKIN::UNC-70(n493)::mKate)* combined with a loss-of-function allele of *tbc-10*, a GAP that inactivates RAB-35 (genotype = *tbc-10; vdSi2*) ^5,17,18^. In these animals the PLM neuron and axon develop normally, but the axon spontaneously breaks and degenerates in adults ^5^ (Fig. 1a-c). From this screen, we identified a recessive mutant, *vd92*, that caused a strong suppression of PLM axon breaks (Fig. 1c). Whole-genome sequencing and direct mapping of *vd92* ^23^ revealed a missense mutation in the *agef-1* locus, which encodes an evolutionarily conserved GEF proposed to activate Arf GTPases implicated in the endocytic-recycling pathway ^21,22,24^. The *vd92* mutant allele carried a substitution of serine by leucine in position 784 (S784L), which corresponds to an auto-inhibitory domain (HDS1) within the AGEF-1a protein, downstream of its catalytic domain (sec7) (Fig. 1d and Fig. S1). To confirm that *vd92* is an allele of *agef-1*, we decided to replicate the identified *agef-1[S784L]* mutation *de novo* using CRISPR-cas9. Indeed, we observed that *agef-1[S784L]* phenocopied *vd92* in suppressing PLM axon breaks when in *tbc-10; vdSi2* background; as predicted, this mutation did not induce PLM axon breaks nor any other detectable phenotype when in a wild-type background (Fig. 1a). Taken together, these results conclusively reveal that *vd92* is an allele of *agef-1*.

**Figure 1.**
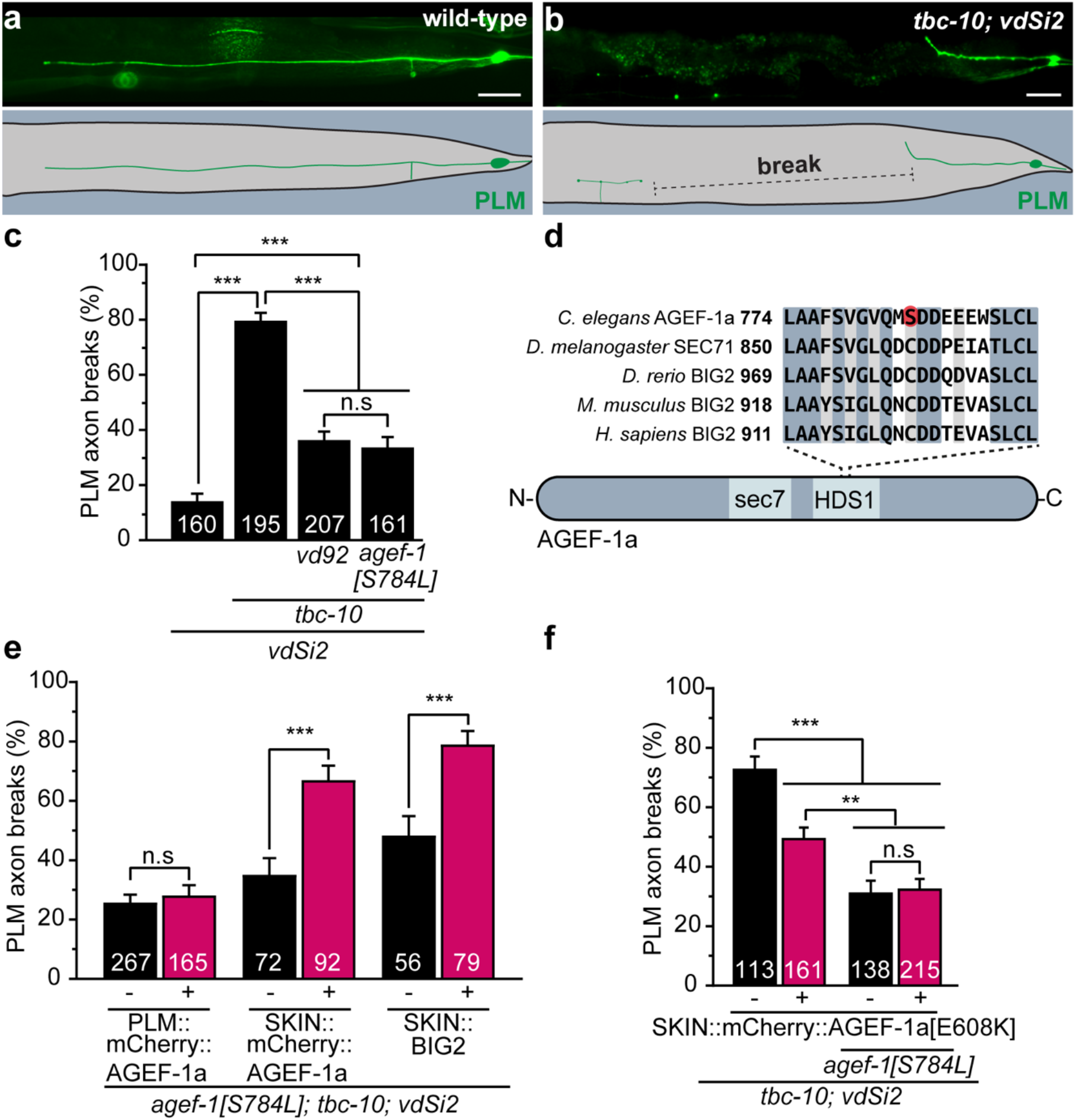
AGEF-1a/BIG2 is conserved and functions non-cell autonomously in the epidermis to modulate axonal integrity. (a-b) Lateral view of the tail of adult animals and schematics of the observed anatomical location. (a) Wild-type, intact PLM axon (Pmec-4::GFP). (b) *vdSi2; tbc-10* sensitized animal, showing a broken PLM axon. Scale bar = 20 µm. (c) Mean penetrance of PLM axon breaks in *vdSi2* single transgenics, *vdSi2; tbc-10* double mutants, *vd92; tbc-10; vdSi2* triple mutants, and CRIPSR-engineered *agef-1[S784L]; tbc-10; vdSi2* triple mutants. (d) Mutated residue within AGEF-1a and its corresponding orthologs. Numbering indicates residue position. Light blue = conserved residues. Grey = equivalent residues. Red = *vd92* mutation site. (e) Mean penetrance of PLM axon breaks in *agef-1[S784L]; tbc-10; vdSi2* triple mutants expressing the *C. elegans* AGEF-1a cDNA selectively within the PLM neurons (PLM = *Pmec-4*) or skin (SKIN = *Pdpy-7*), and human BIG2 cDNA selectively in the skin. (f) Mean penetrance of PLM axon breaks in *vdSi2; tbc-10* and *agef-1[S784L]; tbc-10; vdSi2* animals expressing a dominant-negative *C. elegans* AGEF-1a[E698K] cDNA tissue-specifically in the skin. Bars represent the mean penetrance based on a uniform-prior distribution. Magenta bars represent transgenic animals and black bars their non-transgenic siblings. Standard error of the mean is represented on top of the bars. Sample size represented within the bars. ANOVA used to compare multiple groups. t-test used to compare transgenics vs. non-transgenics. ***, p<0.001; **, p<0.01; n.s = non-significant.

Next, to determine the tissue in which *agef-1* functions to regulate axonal integrity we selectively expressed a wild-type copy of the AGEF-1a cDNA with an N-terminal mCherry tag in either the epidermis (*SKIN::mCherry::AGEF-1a*) or the mechanosensory neurons (*PLM::mCherry::AGEF-1a*), in *vd92; tbc-10; vdSi2* mutant animals. We found that only *SKIN::mCherry::AGEF-1a* was able to rescue the mutant phenotype and restore the rate of PLM axon breaks to background levels (Fig. 1e). Thus, AGEF-1a functions selectively within the epidermis to maintain the PLM axon integrity, in a similar fashion to RAB-35, TBC-10 and UNC-70/β-spectrin ^5^.

*ARFGEF2* is the human ortholog of *agef-1* and encodes for the Brefeldin A-inhibited guanine nucleotide-exchange protein 2 (BIG2); while the human and nematode proteins present a high amino acid sequence conservation, the serine residue mutated in *agef-1[S784L]* is not conserved (Fig. 1d). To determine whether AGEF-1a is functionally conserved, we expressed wild-type human BIG2 cDNA selectively within the epidermis (*SKIN::BIG2*) of *agef-1; tbc-10; vdSi2* animals to determine its capacity to compensate for the loss of AGEF-1a function. Importantly, we found that *SKIN::BIG2* fully restored the rate PLM axon breaks (Fig. 1e), supporting the notion that the molecular function of AGEF-1a in mediating axonal integrity is evolutionarily conserved.

We attempted to generate a full deletion of the *agef-1* locus using CRISPR-cas9, but the resulting deletion was lethal, consistent with previous reports of *agef-1*-mediated viability ^21^. This indicated that the *agef-1[S784L]* is not a null, but likely a loss-of-function allele. Thus, we next decided to investigate the functional consequence of the S784L mutation. Since this mutation falls within an auto-inhibitory domain of AGEF-1a (Fig. 1d), we hypothesized that it may disrupt the GTPase activation function. First, to determine how a disruption to the GTPase activation function would impact axonal integrity, we engineered a dominant negative *agef-1a* cDNA (AGEF-1a[E806K]), which mimics a previously validated mutation in BIG2 that is unable to activate GTPases ^25^. We then over-expressed this dominant negative gene within the epidermis (SKIN::AGEF-1a[E608K]) in *tbc-10; vdSi2* animals, and found that it suppressed the PLM axon breaks, phenocopying *agef-1[S784L]; tbc-10; vdSi2* animals (Fig. 1f). We reasoned that if the *S784L* mutation we isolated causes a similar loss of the GTPase activation function, then SKIN::AGEF-1a[E608K] would not have the capacity to restore the rate of PLM axon breaks to background levels in *agef-1[S784L]; tbc-10; vdSi2* animals. Indeed, we observed that over-expression of SKIN::AGEF-1a[E608K] could not restore the PLM axon breaks in *agef-1[S784L] tbc-10; vdSi2* mutants (Fig. 1f). Thus, we conclude that *agef-1[S784L]* is a partial loss-of-function allele that affects GTPase activation.

In summary, these results indicate that AGEF-1a functions within the epidermis to regulate the integrity of the PLM axon through the activation of GTPases, and that this molecular function is conserved in the human ortholog BIG2.

### AGEF-1a functions upstream of RAB-35

AGEF-1a/BIG2 is known to participate in the endocytic/recycling pathway in multiple cellular contexts by physically binding and activating Arf GTPases, particularly ARF-1.2 and ARF-5 ^21,22,26,27^; however, it is not clear whether Arf GTPases are its exclusive targets ^24,25^. Given the genetic context in which we identified *agef-1*, we hypothesized that AGEF-1a could facilitate the activation of the GTPase RAB-35 to regulate axonal integrity. This could be a direct mechanism that requires AGEF-1a to bind RAB-35, or an indirect mechanism that requires activation of another GTPase, or suite of GTPases, that in turn facilitate activation of RAB-35, as observed in the context of tumor migration ^28^. To address this notion, we took a multifaceted approach.

First, we reasoned that if AGEF-1a functions in RAB-35 activation, it must act upstream of active, GTP-bound RAB-35 (Fig. 2a). To test this possibility, we over-expressed a dominant, constitutively active RAB-35 in the epidermis of *agef-1[S784L]; tbc-10; vdSi2* animals (*SKIN:: RAB-35[Q69L]*). We found that constitutively active RAB-35 fully restored PLM axon breaks in *agef-1[S784L]; tbc-10; vdSi2* compared to their non-transgenic siblings (three independent lines, Fig. 2b), consistent with *agef-1* functioning upstream of active GTP-bound RAB-35.

**Figure 2.**
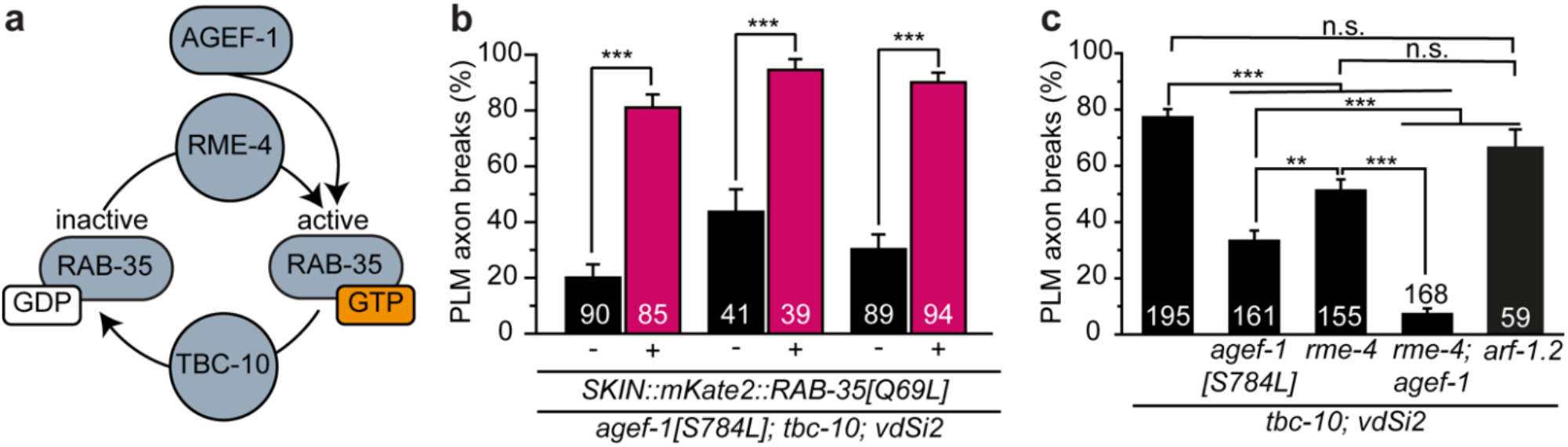
AGEF-1a functions upstream of RAB-35 activity. (a) Schematic of the RAB-35 activation cycle. (b) Mean penetrance of PLM axon breaks in three independent transgenic lines of *agef-1[S784L]; tbc-10; vdSi2* triple mutants expressing a constitutively active mKate2::RAB-35[Q69L] tissue-specifically within the skin (SKIN = *Pdpy-7*). (c) Mean penetrance of PLM axon breaks in *tbc-10*; *vdSi2* sensitized animals, *agef-1[S784L]; tbc-10; vdSi2* triple mutants, *rme-4; tbc-10; vdSi2* triple mutants, and *rme-4; agef-1; tbc-10; vdSi2* quadruple mutants. Bars represent the mean penetrance based on a uniform-prior distribution. Magenta bars represent transgenic animals and black bars their non-transgenic siblings. Standard error of the mean is represented on top of the bars. Sample size represented within the bars. ANOVA used to compare multiple groups. t-test used to compare transgenics vs. non-transgenics. ***, p<0.001; **, p<0.01; n.s = non-significant.

Next, we decided to test whether AGEF-1 functions in synergy with other known activators of RAB-35 (Fig. 2a). To test this idea, we first determined the genetic interaction between *agef-1* and *rme-4*, a known RAB-35 GEF that partially modulates axonal integrity ^5,18^. In *tbc-10; vdSi2* animals, the null allele *rme-4(b1001)* ^18^ partially suppresses PLM axon breaks, although less effectively than *agef-1[S784L]* (Fig. 2c and ^5^). In contrast, *rme-4; agef-1; tbc-10; vdSi2* animals displayed a full suppression of PLM axon breaks (Fig. 2c), supporting the notion that these molecules act in synergy.

Expanding on this concept, given that AGEF-1a/BIG2 is known to activate ARF-1/Arf1 and ARF-5/Arf3 in the context of endocytic recycling ^19^, we decided to test whether AGEF-1a facilitates RAB-35 activation through these specific GTPases. We reasoned that if ARF-1.2/5 were involved, their loss would suppress PLM breaks in *tbc-10; vdSi2* animals. To test this possibility, we used CRISPRcas9 to knock-out *arf-1*.*2* and *arf-5* in *tbc-10; vdSi2* mutant animals. *arf-5* deletion led to fully penetrant larval lethality, precluding isolation of mutants and analysis of this gene. Surprisingly, we observed that lack of *arf-1*.*2* did not affect PLM axon breaks in the *tbc-10; vdSi2* animals (Fig. 2c). These results indicate that AGEF-1a facilitates RAB-35-activation independently of ARF-1.2.

Taken together, our data support a model where AGEF-1a facilitates RAB-35 activation in synergy with RME-4. Furthermore, since AGEF-1a is functioning independently of its classical ARF-1.2 in this context, it is possible that it may activate RAB-35 directly.

### AGEF-1a interacts with RAB-35 in the epidermis

Given our genetic data, we hypothesized that AGEF-1a could be interacting directly with RAB-35. Thus, we investigated the subcellular localization of AGEF-1a and RAB-35 within the epidermis. We leveraged a split-fluorophore strategy to visualize endogenous AGEF-1a within the epidermis by using CRISPR-cas9 to fuse 7 tandem repeats of GFP^11^ with the C-terminus of endogenous AGEF-1a (AGEF-1a::GFP^11^) ^29^. We then expressed GFP^1-10^ under an epidermal-specific promoter (SKIN::GFP^1-10^) to reconstitute full-length GFP selectively within the epidermis (epidermal AGEF-1a::GFP_x7_). Simultaneously, we visualized RAB-35 by expressing a functional wild-type RAB-35 carrying an N-terminal mKate tag under the control of an epidermal specific promoter (SKIN::mKate::RAB-35) ^5^, as well as carrying a null allele of *rab-35(b1013)* to minimize any over-expression artifacts. We then used super-resolution microscopy to observe the colocalization of both these molecules *in vivo* in the last larval stage (L4; Fig. 3a). Epidermal AGEF-1a::GFP_x7_ localized to stable small puncta throughout the cytoplasm and membrane of the epidermis (Fig. 3b-d). Qualitatively, we did not observe any enrichment of epidermal AGEF-1a::GFP_x7_ in the vicinity of the PLM neuron axon (Fig. 3b, c). mKate::RAB-35, however, localized to punctate, and vesicle-like structures with a clear luminal space within the epidermis and was notably enriched at the membrane near the epidermal furrow that surrounds the PLM axon (Fig. 3b-d, and ^5^). While both molecules do not fully overlap, epidermal AGEF-1a::GFP11_x7_ puncta were frequently found decorating a subset of larger mKate::RAB-35 structures; 3D reconstructions revealed that AGEF-1a::GFP11_x7_ closely interacts with the edge of the mKate::RAB-35 structures (Fig. 3d, e).

**Figure 3.**
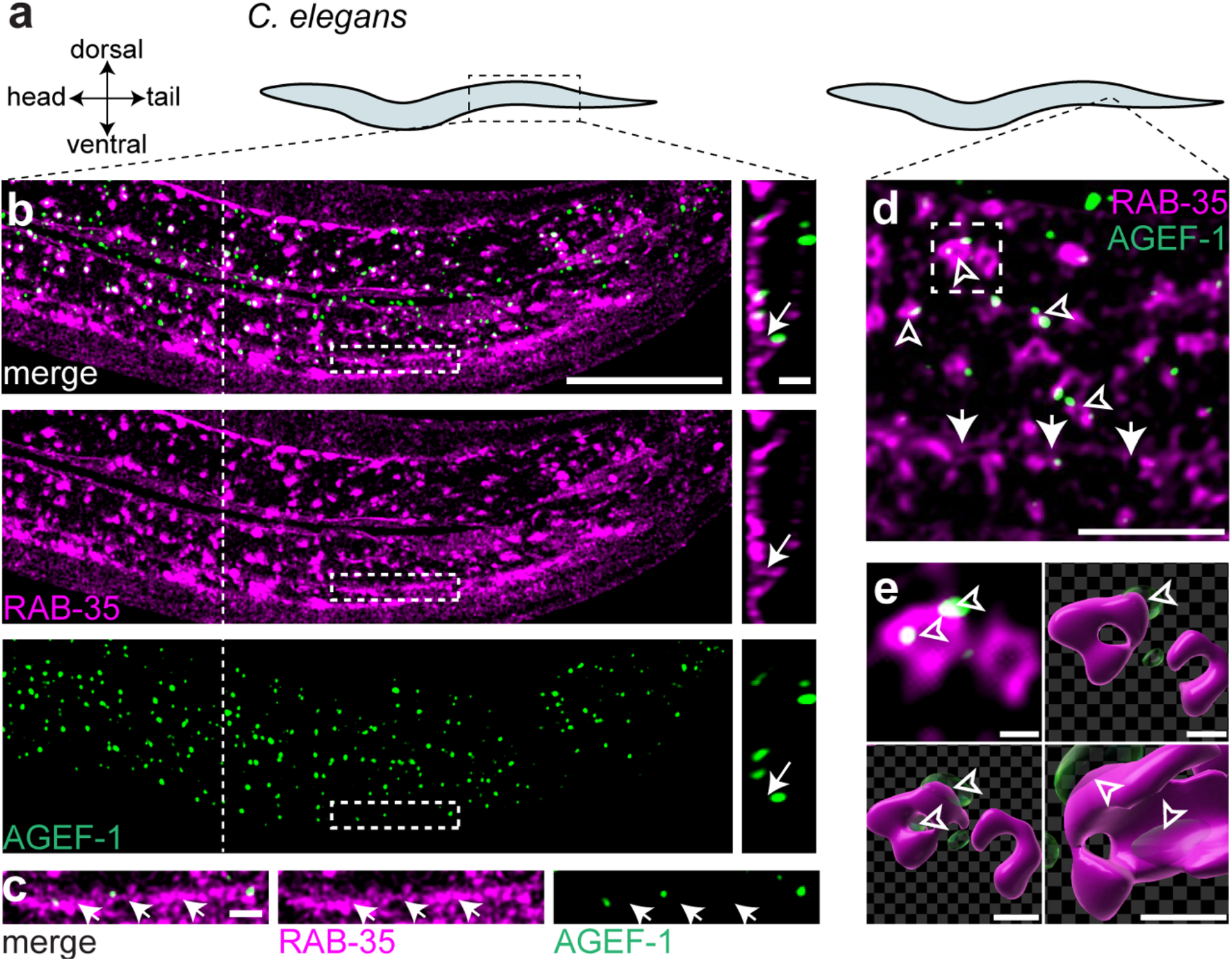
AGEF-1a interacts with RAB-35 *in vivo* within the epidermis. (a) Schematic of a lateral view of the animal highlighting the anatomical region shown in the panels b and d. (b) Representative deconvolved confocal microscopy of the epidermis of a *rab-35(b1013)* animal expressing wild-type mKate::RAB-35 selectively within the skin and endogenous epidermal AGEF-1::GFPx7 (AGEF-1::GFP11×7 + SKIN::spGFP1-10). Right panel is an orthogonal perspective of the region highlighted by the dashed line. Left panel scale bar = 20 µm. Right panel scale bar = 2 µm. (c) Magnified view of the dashed rectangles in panel b. Scale bar = 2 µm. (d) Representative deconvolved array detector super-resolution microscopy of the epidermis of a *rab-35(b1013)* animal expressing wild-type mKate::RAB-35 tissue-specifically within the skin and endogenous epidermal AGEF-1::GFPx7. Scale bar = 10 µm. (e) 3D surface rendering of the boxed region in d in different perspectives. Scale bar = 0.5 µm. Arrows highlight the epidermal furrow. Arrowheads highlight RAB-35 structures decorated with AGEF-1.

Collectively, these data demonstrate that AGEF-1a interacts with a subset of RAB-35 molecules in the epidermis, further supporting the model in which AGEF-1a directly activates RAB-35.

### AGEF-1 alters neuron-epidermal attachments

Correct development and maintenance of uniform neuron-epidermal attachments is essential to shield the PLM axon from breakage induced by the mechanical stress of body movement ^1,5^. RAB-35 activity fine-tunes the resiliency of these specialized attachments in synergy with epidermal β-spectrin, with the appearance of discontinuous neuron-epidermal attachments after movement being predictive of axonal damage ^5^. We next asked whether AGEF-1a also affects neuron-epidermal attachments via RAB-35. If AGEF-a functions upstream of RAB-35 to facilitate its activation, we predicted that AGEF-1a loss-of-function would protect neurons by suppressing the neuron-epidermal attachment defects present in *tbc-10; vdSi2* animals. To investigate this aspect, we observed the localization of a key epidermal attachment molecule, LET-805, that is part of the neuron-epidermal attachment complex ^1,15,30^. To visualize this molecule, we engineered transgenic animals in which the endogenous *let-805* locus is fused with the sequence encoding the fluorophore wrmScarlet ^1,5^. We then analyzed the localization of LET-805::wrmScarlet in *agef-1[S784L]; tbc-10; vdSi2* animals in comparison with *tbc-10; vdSi2* during the last larval stage (L4), and after the appearance of PLM axon breaks (1DOA) (Fig. 4a, b). We found that at the L4 stage, *agef-1[S784L]; tbc-10; vdSi2* did not significantly differ from *tbc-10; vdSi2* (Fig. 4c), and in adults all *tbc-10; vdSi2* animals presented a discontinuous LET-805::wrmScarlet localization along the PLM axon vicinity, with most of them accompanied by a PLM break (Fig. 4b, c).

**Figure 4.**
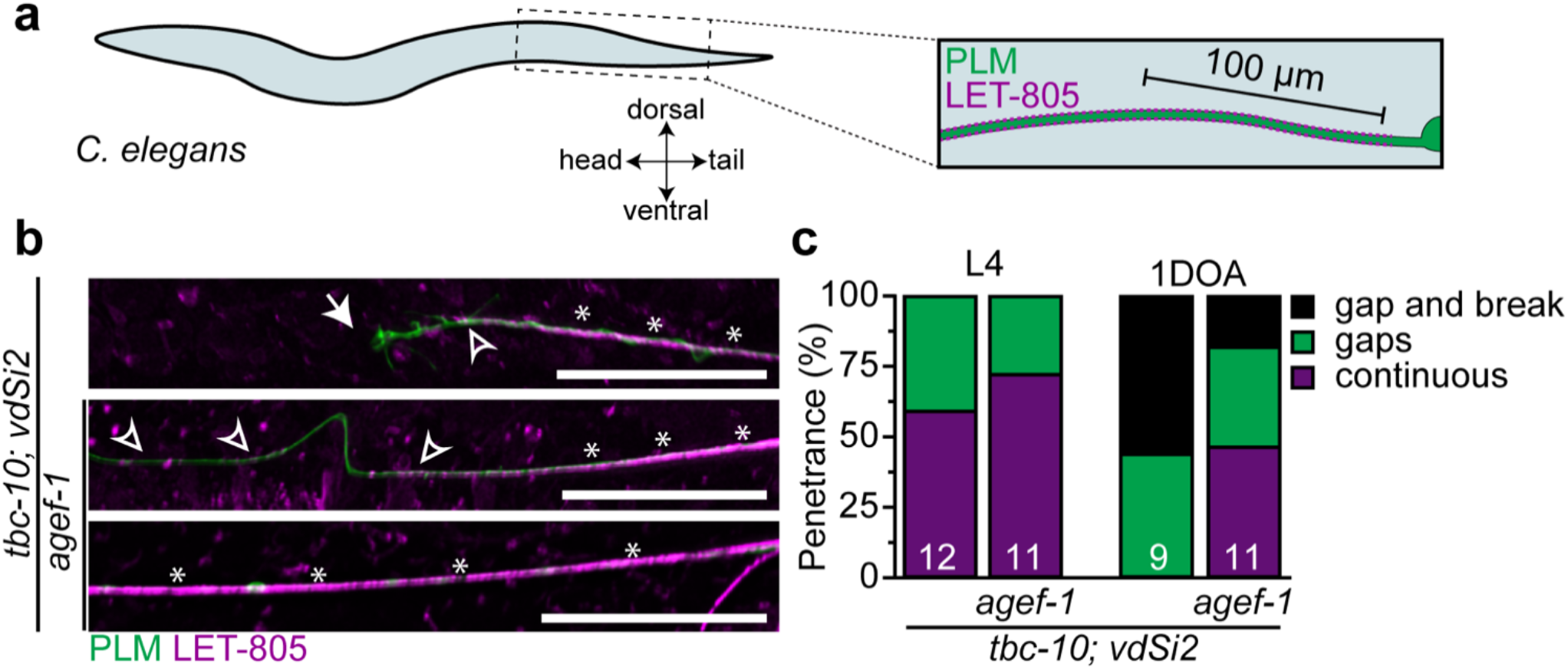
AGEF-1a alters neuron-epidermal attachment. (a) Schematic a lateral view the animal highlighting the anatomical region analysed. (b) Representative images of the quantified phenotypes, showing a lateral view of the PLM axon and endogenous LET-805::wrmScarlet. Top panel represents a *tbc-10; vdSi2* animal with the ‘gap a break’ phenotype. Middle panel represents an *agef-1; tbc-10; vdSi2* triple mutant animal with the ‘gaps’ phenotype. Bottom panel represents an *agef-1; tbc-10; vdSi2* triple mutant with the ‘continuous’ phenotype. Arrows highlight a PLM axon break. Arrowheads highlight naked PLM axons. Asterisks highlight regions of consistent neuron-epidermal attachment. Scale bar = 20 µm. (c) Quantification of the phenotypes observed in panel b showing the proportion of *tbc-10; vdSi2* sensitized animals and *agef-1; tbc-10; vdSi2* triple mutants at the last larval stage (L4) and 1 day-old adult (1DOA) stage. Sample size represented within the bars.

In stark contrast, approximately half of the *agef-1[S784L]; tbc-10; vdSi2* animals retained a continuous LET-805::wrmScarlet localization (Fig. 4b, c). Of the adult *agef-1[S784L]; tbc-10; vdSi2* animals that displayed gaps in LET-805::wrmScarlet localization near the PLM axon region, ∼67% displayed an intact axon (however, notably buckled), whereas the remaining ∼33% were broken (Fig. 4b, c).

Together, these results conclusively demonstrate that AGEF-1a loss-of-function suppresses the neuron-epidermal attachments defect present in *tbc-10; vdSi2* animals, protecting the associated sensory axons from movement-induced damage.

## Discussion

By leveraging the power of unbiased forward genetics, we reveal that AGEF-1a regulates the axonal integrity of mechanosensory neurons in *C. elegans* by interacting with, and facilitating the activation of, RAB-35 in the epidermis to modulate the stability of neuron-epidermal attachments. Furthermore, we demonstrate the evolutionary conservation of this molecular function by showing that the human ortholog, BIG2, can replace AGEF-1 in the context of axonal maintenance. Importantly, our results highlight a trans-tissue maintenance mechanism whereby the integrity of mechanosensory neurons are regulated by components functioning non-cell autonomously within the epidermis. Finally, our results support a model where uniform neuron-epidermal attachments are tightly regulated to preserve the integrity of axons.

Mutations in BIG2 are associated with the development of familial microcephaly, and disruption of this molecule causes neuronal migration defects in culture models ^31–33^. The mechanism of action of BIG2 in neuronal health is unknown and, in general, research in this area focuses on neuron-intrinsic functions of this molecule. Here we show that AGEF-1a/BIG2 can function non-cell-autonomously outside the nervous system to regulate axonal integrity, and that this molecular role is conserved. Future investigations of BIG2-associated neurological disorders should consider a non-neuronal function, and hyper-activity of BIG2 as a driver of neuropathology. Moreover, given that we observe a detrimental function of AGEF-1a/BIG2 in the context of a spectrin disruption, we propose that mutations in BIG2 could be disease-modifiers in patients with spectrinopathies.

To our surprise, AGEF-1a functions independently of its classical target GTPase ARF-1.2 in the context of axonal maintenance, but still dependent on its GTPase activating function. This suggests that either AGEF-1a is directly mediating RAB-35 GDP release and GTP binding, or the existence of an unknown intermediate GTPase. Others have shown that Arf5 allosterically activates DENND1 (ortholog of RME-4) GEF activity towards Rab35 in mice glioblastoma derived cells ^28^. Interestingly, BIG2 expression induces a small increase in Arf5-GTP levels in HeLa cells ^19^. Moreover, nematode ARF-1.2, ARF-5 and human Arf5 share more than 80% sequence identity, suggesting a possible functional redundancy and conservation. Given these results, and our observation that AGEF-1a functions in synergy with RME-4 and upstream of RAB-35 activity, it is possible that ARF-5 is a target of AGEF-1a. Addressing whether these four molecules form a complex *in vivo* will be key to answering this question.

How do AGEF-1a, RAB-35, and UNC-70 affect neuron-epidermal attachment? AGEF-1a is a known modulator of the basolateral polarity of the epidermal growth factor LET-23 and protein secretion ^21^. Furthermore, it is implicated in the regulation of endosome/lysosome fusion/fission as well as endocytic transport ^22^. RAB-35 and UNC-70 have also been described to be involved in different steps of endocytosis and membrane recycling, and their exact function in this processes seems complex ^10,18,34–36^. In line with these findings, our work supports the idea that UNC-70 stabilizes adhesive membrane domains, whereas RAB-35 and AGEF-1a activity further destabilize these attachments in the absence of UNC-70. These notions support the concept that membrane recycling events within the epidermal membrane closest to an innervating neurite are critical for the maintenance of axonal integrity. Consequently, a trans-tissue coordination of membrane domain organization is required to support organismal health, especially between tissues that are under constant mechanical challenge.

## Methods

### Strains

Nematode cultures were grown in nematode growth medium (NGM) plates seeded with OP50 *Escherichia coli* and maintained at 20 °C following standard methods ^37^. Animals were visually selected at the last larval stage on a stereomicroscope and then scored for PLM axon breaks 48 hours later as 2-day-old adults. Mutant animals of genotype *tbc-10(vd31); vdSi2[Pdpy-7::UNC-70(n493)::mKate2]; zdIs5[Pmec-4::GFP]* were generated previously ^5^. Strains used in this work are listed in Table S1 and S2.

### Mutant strains isolation

Animals of genotype *vdSi2[Pdpy-7::UNC-70(n493)::mKate2]; tbc-10(vd31); zdIs5* were mutagenized using 50 mM ethyl methanesulfonate (EMS, Sigma) for 4 hours. *vd92* mutants were isolated in a clonal F2 progeny screen and then backcrossed three times with the original sensitized strain QH7929 [*tbc-10(vd31); vdSi2[Pdpy-7::unc-70(n493)::mKate2]* before whole-genome sequencing was conducted (GeneWiz). Strains QH8151 [*vd92; tbc-10(vd31); vdSi2[Pdpy-7::UNC-70(n493)::mKate2]; zdIs5[Pmec-4::GFP]*], QH7929 [*vdSi2[Pdpy-7::UNC-70(n493)::mKate2]; tbc-10(vd31); zdIs5[Pmec-4::GFP]*], CZ10175 [*zdIs5[Pmec-4::GFP]*] and the N2 Bristol strain were whole-genome sequenced at a mean depth of ≥ 19. Genome mapping and calling of variants were performed by GeneWiz. Background variants subtraction and candidate mutation filtering and mapping was performed using a customized Python script (https://github.com/IgorBonaP/variations_mapper) following a strategy of direct mapping ^23^.

### Genetic engineering

Knock-ins, single base pair changes, and deletions were made with CRISPR-cas9 using a co-CRISPR strategy ^38^. An equimolar mix of ALT-R Cas9, tracRNA and ALT-R crRNA (Integrated DNA Technologies) at a final concentration of 1.525 μM plus single-stranded oligonucleotides (Integrated DNA Technologies) repair templates at a final concentration of 1.6 μM, or PCR products in a concentration range of 0.3-0.5 μM, were diluted in ultrapure water and microinjected into the gonads of adult animals. All repair templates were designed with 75 base pair homology to flanking genomic regions. PCR-repair templates were amplified using ultramer oligonucleotides (Integrated DNA Technologies). For deletions, no repair templates were used. Sequences of all crRNAs and oligonucleotides used are listed in Tables S3. Final engineered lines were genotyped by PCR and Sanger sequencing.

### Transgenics

Transgenic strains carrying extrachromosomal arrays were generated by microinjection into the germline using standard methods ^39^. All constructs were made using the Gibson isothermal assembly method ^40,41^ except Pdpy-7::mKate::AGEF-1a[608K] and Pdpy-7::mKate::RAB-35[Q79L], which were generated using the quick-change mutagenesis II kit (Agilent). DNA fragments for assembly were generated by PCR. pSM vectors were utilized as the backbone for assembly. The sequence of all constructs was validated by Sanger sequencing. The sequence of all constructs is listed in Table S2. AGEF-1a and BIG2 cDNA were obtained from Genescript based on the sequence of WormBaseID:Y6B3A.1a ^42^. AGEF-1a[E608K] sequence was determined by protein alignment of AGEF-1a and BIG2[E738K] ^25^ then generated using construct Pdpy-7::mKate::AGEF-1a as template.

### Microscopy

Animals were inspected by mounting on 4% agar pads after anesthesia with 0.05% tetramisole hydrochloride in M9 buffer. Fluorescence microscopy was performed using an upright Zeiss AxioImager A1 microscope equipped with a Photometrics Cool Snap HQ2 camera. Images acquired in Metamorph®. Confocal imaging was performed using a spinning disk confocal microscope (Marianas, Intelligent Imaging Innovations), equipped with a confocal scanner unit (CSU-W1, Yokogawa Electric Co.) built around an Axio Observer body (Z1, Carl Zeiss AG) and fitted with an sCMOS camera (ORCA-Flash4.0 V2, Hamamatsu Photonics) and SlideBook v6.0 software (3i) using a 63×/1.4 NA oil-immersion objective with sampling intervals x,y = 99 nm and z = 130 nm. Image acquisition was performed using SlideBook 6.0 (3I, Inc). Detector-array super resolution imaging was performed using a Zeiss C Plan-Apochromat 63x/1.4 NA oil-immersion objective on a confocal/two-photon laser-scanning microscope (LSM 980 NLO Airyscan 2, Carl Zeiss GmbH) built around an Axio Observer 7 body and equipped with an Airyscan 2 super-resolution detector, a 34-channel spectral photomultiplier tube (PMT) array, two internal GaAsP PMTs, a transmission PMT, and two external GaAsP PMTs for non-descanned detection in two-photon microscopy. Both confocal and detector-array imaging were done in the Queensland Brain Institute’s Advanced Microscopy Facility. Confocal microscope acquisitions were deconvolved using Huygens Professional version Huygens Professional v18.04 (Scientific Volume Imaging, The Netherlands, http://svi.nl) run on a GPU-accelerated computer (3x NVIDIA® Tesla® V100), using the CMLE algorithm with SNR:20 and 40 iterations. Detector-array microscope acquisitions were deconvolved using the GMLE algorithm, with SNR:8.31, Acuity: 20 and 10 iterations. Microinjections were performed using standard methods ^39^, on either an inverted Zeiss AxioObserver microscope equipped with differential interference contrast, a Narishige needle holder, and a Tritech Research injection system.

### Image analysis and panel preparation

All images were analyzed in FIJI ^43^. LET-805::wrmScarlet localization scoring was done using a custom FIJI macro to segment the analysis region with the aid of the plugin SNT ^44^ to trace the axon and then generate a line profile of the LET-805::wrmScarlet channel in that area. A custom Python script was made to analyze the first 100 µm of the PLM axon line profiles. Using this data as input, the program then scored animals with gaps in LET-805::wrmScarlet larger than ∼5 µm as ‘gaps’, animals with continuous LET-805::wrmScarlet as ‘continuous’ and animals with a PLM break as ‘gap and break’. A gap was defined as a bin of ∼5 µm in length where the sum of its signal is less than 21% of the maximum possible signal in such a bin. Figure panels were prepared using FigureJ ^45^ and Adobe Illustrator®.

### Protein sequence alignment and modeling

AGEF-1a amino acid sequence was aligned to its closest orthologs in *Danio rerio, Drosophila melanogaster, Mus musculus* and *Homo sapiens* using the alignment tool in www.uniprot.org ^46^. Respective uniprot ID for each molecule: G5EFH7_CAEEL, E7FCG1_DANRE, Q9VJW1_DROME, BIG2_MOUSE, BIG2_HUMAN. Uniprot IDs used for sequence alignment of nematode ARF-1.2, ARF-5 and human Arf5: ARF12_CAEEL, G5EFK4_CAEEL, ARF5_HUMAN. 3d model generated using AlphaFold^47^.

## Statistical analysis

Statistical analysis and plotting were performed using GraphPad Prism version 11 for Windows (GraphPad Software, La Jolla California USA, www.graphpad.com). One-way ANOVA followed by Tukey’s multiple comparisons test was used to compare the mean of multiple groups. *t-*test was used to compare pairs.

## Author contribution

SC and IBP performed experiments, collected, and analyzed data. SC, IBP, and MAH conceptualized and designed experiments, discussed results, interpreted data, and wrote the manuscript.

## Funding

NHMRC-ARC Dementia Development Fellowship APP1108489, and Ideas Grant GNT2010532 (SC)

NHMRC Investigator Grant APP1197860 and Project Grant APP1129546 (MAH) Alastair Rushworth Research Fund (IBP)

ARC LIEF grant LE130100078 (QBI Advanced Microscopy Facility)

## Acknowledgments

We thank R. Amor, A. Thompson, and A. Gaudin for support with microscopy; colleagues in the Hilliard and Coakley laboratories for insightful discussions and comments. Some strains were provided by the CGC, which is funded by NIH Office of Research Infrastructure Programs (P40 OD010440), and the International *C. elegans* Gene Knockout Consortium.

## Notes

### Competing Interest Statement

The authors have declared no competing interest.

