## Supplementary Material for "A conserved Arf-GEF modulates axonal integrity through RAB-35 by altering neuron-epidermal attachment"

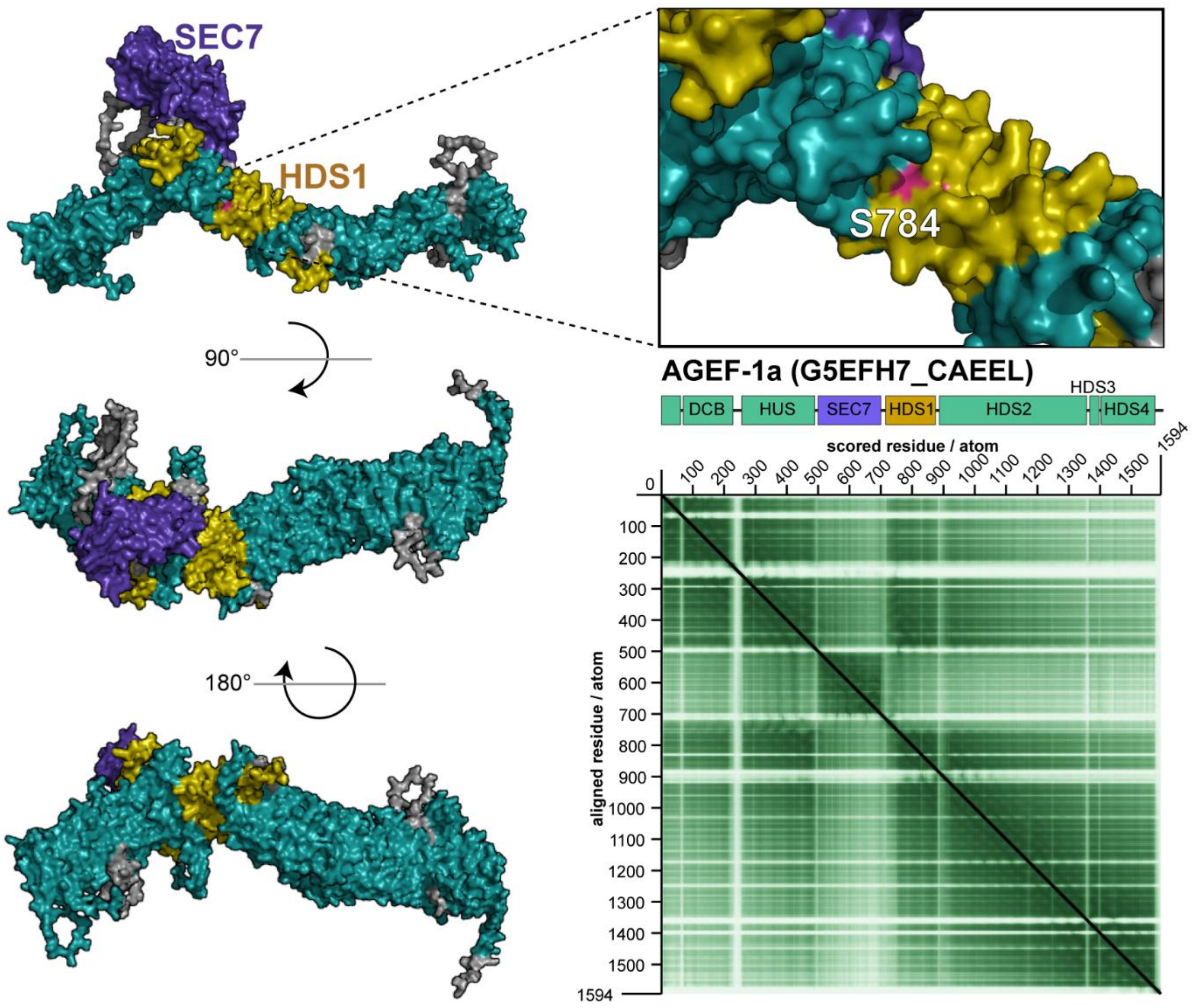


**Figure S1. AlphaFold model of AGEF-1a.** 3d structural model of AGEf-1a highlighting the location of the catalytic site (SEC7 domain) and the *vd92* allele causative mutation (S784) and the position alignment error plot or the predicted structure.

| Strain | Genotype | Stage | PLM break | n | Figure # |
|---|---|---|---|---|---|
| QH7929 | *vdSi2(Pdpy-7::UNC-70(n493)::mKate2) X; tbc-10(vd31) III; zdIs5(Pmec-4::GFP) I* | 2DOA | 156 | 195 | *1a* |
| QH6753 | *vdSi2[Pdpy-7::UNC-70(n493)::mKate2] X; zdIs5(Pmec-4::GFP) I* | 2DOA | 22 | 160 | *1a* |
| QH8151 | *agef-1(vd92*[S784L]) I; vdSi2(Pdpy-7::UNC-70(n493)::mKate2) X; tbc-10(vd31) III; zdIs5(Pmec-4::GFP) I* | 2DOA | 75 | 207 | *1a* |
| QH8738 | *agef-1(vd123*[S784L]) I; vdSi2(Pdpy-7::UNC-70(n493)::mKate2) X; tbc-10(vd31) III; zdIs5(Pmec-4::GFP) I* | 2DOA | 54 | 161 | *1a, 2c* |
| QH9584 | *rme-4(b1001) vdSi2(Pdpy-7::UNC-70(n493)::mKate2) X; tbc-10(vd31) III; zdIs5(Pmec-4::GFP) I* | 2DOA | 80 | 155 | *2c* |
| QH9583 | *agef-1(vd123*[S784L]) I; rme-4(b1001) vdSi2(Pdpy-7::UNC-70(n493)::mKate2) X; tbc-10(vd31) III; zdIs5(Pmec-4::GFP) I* | 2DOA | 12 | 168 | *2c* |
| QH220 | *arf-1(vd220) III; vdSi2(Pdpy-7::UNC-70(n493)::mKate2) X; tbc-10(vd31) III; zdIs5(Pmec-4::GFP) I* | 2DOA | 40 | 59 | *2c* |
| QH9616 | *agef-1(vd92*[S784L]) I; vdSi2(vd187*[Pdpy-7::UNC-70(n493)]) X; let-805(syb380[let-805::wrmScarlet] tbc-10(vd31) III; zdIs5(Pmec-4::GFP) I* | L4, 2DOA | -- | -- | *4b* |
| QH9620 | *vdSi2(vd187*[Pdpy-7::UNC-70(n493)]) X; let-805(syb380[let-805::wrmScarlet] tbc-10(vd31) III; zdIs5(Pmec-4::GFP) I* | L4, 2DOA | -- | -- | *4b* |

**Table S1. Stable lines utilized in this study.**

| Strain | Genotype | Stage | Transgenic PLM break | Transgenic n | Non-transgenic PLM break | Non-transgenic n | Figure # | |
|---|---|---|---|---|---|---|---|---|
| QH8855 | vdEx2527[Pdpy-7::mCherry::AGEF-1a (1.25 ng/μL)]; tbc-10 (vd30) III; vdSi2[Pdpy-7::UNC-70(n493)::mKate2] X; agef-1(vd92) zdIs5(Pmec-4::GFP) I | 2DOA | 62 | 92 | 25 | 72 | 1e | |
| QH8942 | vdEx2525[Pdpy-7::BIG2 (2.5ng/ul)]; tbc-10 (vd30) III; vdSi2[Pdpy-7::UNC-70(n493)::mKate2] X; agef-1(vd92) zdIs5(Pmec-4::GFP) I | 2DOA | 63 | 79 | 27 | 56 | 1e | |
| QH9838 | vdEx2871[Pmec-4::mCherry::AGEF-1a (10 ng/μL)]; tbc-10 (vd30) III; vdSi2[Pdpy-7::UNC-70(n493)::mKate2] X; agef-1(vd92) zdIs5(Pmec-4::GFP) I | 2DOA | 16 | 81 | 20 | 72 | 1e | |
| QH9836 | vdEx2869[Pmec-4::mCherry::AGEF-1a (10 ng/μL)]; tbc-10 (vd30) III; vdSi2[Pdpy-7::UNC-70(n493)::mKate2] X; agef-1(vd92) zdIs5(Pmec-4::GFP) I | 2DOA | 29 | 88 | 5 | 36 | 1e | Pooled |
| QH9837 | vdEx2870[Pmec-4::mCherry::AGEF-1a (10 ng/μL)]; tbc-10 (vd30) III; vdSi2[Pdpy-7::UNC-70(n493)::mKate2] X; agef-1(vd92) zdIs5(Pmec-4::GFP) I | 2DOA | 23 | 98 | 21 | 57 | 1e | |
| QH9984 | vdEx2888[Pdpy-7::mCherry::AGEF-1[E608K] (2 ng/μL)]; vdSi2[Pdpy-7::UNC-70(n493)::mKate2] X; tbc-10(vd31) III; zdIs5(Pmec-4::GFP) I | 2DOA | 80 | 161 | 83 | 113 | 1f | |
| QH10092 | vdEx2922[Pdpy-7::mKate2::RAB-36[Q69L]]; agef-1(vd123) I; vdSi2[Pdpy-7::UNC-70(n493)::mKate2] X; tbc-10(vd31) III; zdIs5(Pmec-4::GFP) I | 2DOA | 70 | 85 | 18 | 90 | 1f | |
| QH10033 | vdEx2888[Pdpy-7::mCherry::AGEF-1[E608K] (2 ng/μL)]; vdSi2[Pdpy-7::UNC-70(n493)::mKate2] X; tbc-10(vd31) III; agef-1(vd123) zdIs5(Pmec-4::GFP) I | 2DOA | 70 | 215 | 43 | 138 | 1f | |
| QH10092 | vdEx2922[Pdpy-7::mKate2::RAB-35[Q69L] (2 ng/μL)]; agef-1(vd123*[S784L]) I; vdSi2[Pdpy-7::UNC-70(n493)::mKate2] X; tbc-10(vd31) III; zdIs5(Pmec-4::GFP) I | 2DOA | 70 | 85 | 18 | 90 | 2b | |
| QH10093 | vdEx2923[Pdpy-7::mKate2::RAB-35[Q69L] (2 ng/μL)]; agef-1(vd123*[S784L]) I; vdSi2[Pdpy-7::UNC-70(n493)::mKate2] X; tbc-10(vd31) III; zdIs5(Pmec-4::GFP) I | 2DOA | 38 | 39 | 18 | 41 | 2b | |
| QH10094 | vdEx2924[Pdpy-7::mKate2::RAB-35[Q69L] (2 ng/μL)]; agef-1(vd123*[S784L]) I; vdSi2[Pdpy-7::UNC-70(n493)::mKate2] X; tbc-10(vd31) III; zdIs5(Pmec-4::GFP) I | 2DOA | 86 | 94 | 27 | 89 | 2b | |
| QH10118 | rab-35(b1013) III; agef-1(vd124[agef-1::GFP11x7]) I; vdEx1571[Pdpy-7::mKate2::RAB-35 0.2 ng/μl]; vdEx2432[Pdpy-7::GFP1-10 2.5 ng/μl] | L4 | -- | -- | -- | -- | 3b-e | |

**Table S2. Semi-stable transgenic lines utilized in this study.**

| ID | Type | Sequence | Description |
|---|---|---|---|
| IBP253 | Oligo | *caatgttcaaaatctgctggacCccatgccttgccgcattctcagtcggtgtacaaatgtTggatgatgaagaagaatggtctctttgtCtACgaggGttCcgtctAggagtacgtgccg* | Repair template to generate *agef-1[S784L]* |
| G15 | crRNA | *TGTTTGAGAGGTTTTCGTCT* | crRNA to target *agef-1* near the predicted *vd92* mutation. Used to make *agef-1(vd123*[S784L])* |
| G16 | crRNA | *GAGAAATTAATCAGAGCTGT* | crRNA to target *agef-1* c-terminus. Used to make *agef-1::GFP11x7* and to delete *agef-1* |
| IBP251 | Oligo | *cgaataatctaatttaaatacaaaagagccacacaaaaatgagaaattaatcagagctgtgTTgagctagctTCACATGAATTTGCCGCCGGTGATACCGG* | Primer to amplify GFP11x7 with homology to AGEF-1c terminus end. |
| IBP252 | Oligo | *CAAACGAAATCACCAACAACTGTTCAAGAAGCGCTAGAAGAATCTCAGAAATCCCATATTTCGGAAAATTTCACTATGCGTGACCACATGGTCCTTCATGAG* | Primer to amplify GFP11x7 with homology to AGEF-1c terminus end. |
| G46 | crRNA | *aacatgagcgaattgtcaaa* | crRNA to target *agef-1* n-terminus. Used to delete *agef-1.* |
| G34 | crRNA | *tTTAAGATCTATTCTTGAGC* | crRNA to target *arf-1.2* c-terminus. Used to delete arf-1.2 |
| G41 | crRNA | *TATTAATCAATTTATAGAAT* | crRNA to target *arf-1.2* n-terminus. Used to delete arf-1.2 |
| G37 | crRNA | *GACATGTAAGTTGTGCAATT* | crRNA to target *arf-5* c-terminus. Used to delete *arf-5* |
| G45 | crRNA | *ggagattgttaaacccata* | crRNA to target *arf-5* n-terminus. Used to delete arf-5 |

**Table S3. Oligos and crRNAs utilized in this study.**
